# Ectopic DNA integration and marker-free CRISPR/Cas9 strategies for the halotolerant black yeast *Hortaea werneckii*

**DOI:** 10.1101/2024.03.06.583723

**Authors:** Yainitza Hernandez-Rodriguez, A. Makenzie Bullard, Rebecca J. Busch, Aidan Marshall, José M. Vargas-Muñiz

## Abstract

*Hortaea werneckii* is a halotolerant black yeast commonly found in hypersaline environments. This yeast is also the causative agent of tinea nigra, a superficial mycosis of the palm of the hand and soles of the feet of humans. In addition to their remarkable halotolerance, this black yeast exhibits an unconventional cell division cycle, alternating between fission and budding cell division. Cell density and the salt concentration in their environment regulate which cell division cycle *H. werneckii* uses. Although *H. werneckii* have been extensively studied due to their unique physiology and cell biology, deciphering the underlying mechanisms behind these remarkable phenotypes has been limited due to the lack of genetic tools available. Here, we report a new ectopic integration protocol for *H. werneckii* using PEG-CaCl_2_ mediated protoplast transformation. This approach relies on a drug (hygromycin B) resistance gene to select for successful integration of the genetic construct. The same construct was used to express cytosolic green fluorescent protein. Finally, we developed a marker-free CRISPR/Cas9 protocol for targeted gene deletion using the melanin synthesis pathway as a visual reporter of successful transformation. These transformation strategies will allow testing hypotheses related to *H. werneckii* cell biology and physiology.

**Importance:** *Hortaea werneckii* is a remarkable yeast capable of growing in high salt concentration, and its cell division cycle alternates between fission-like and budding. For these unique attributes, *H. werneckii* has gathered interest in a research program studying extremophile fungi and cell division. Most of our understanding of *H. werneckii* biology comes from genomic analyses, usage of drugs to target a particular pathway or heterologous expression of its gene in *S. cerevisiae*. Nonetheless, *H. werneckii* has remained genetically intractable. Here, we report on two strategies to transform *H. werneckii*: ectopic integration of a plasmid and gene deletion using CRISPR/Cas9. These approaches will be fundamental to expanding the experimental techniques available to study *H. werneckii*, including live cell imaging of cellular processes and reverse genetic approaches.

## Introduction

*Hortaea werneckii* (*Dothideales, Ascomycota*) is a black yeast of particular interest due to its ability to grow in high salinity (1). This black yeast is commonly isolated from hypersaline environments, including seawater and solar salterns, and can grow in saturated sodium chloride solution. Not only does this yeast display a remarkable halotolerance, but it also exhibits an unconventional cell division cycle (2). *H. werneckii* first grows in a pattern similar to that of fission yeast. After the first septation, it switches to budding from the poles. This division pattern and cell morphology depend on the growth environment and cell density (3, 4). *H. werneckii* is also of clinical interest due to its ability to cause a superficial mycosis of the hands and feet known as tinea nigra (5– 7). Tinea nigra frequently occurs in countries located in the tropics (6). On rare occasions, *H. werneckii* can cause a systemic infection known as disseminated phaeohyphomycosis (7). *H. werneckii* isolates exhibit great phenotypic diversity, including drug susceptibility and pathogenicity variation (3, 8, 9). This phenotypic diversity also correlates with the genetic diversity of *H. werneckii. H. werneckii* isolates are intraspecific hybrids of two divergent isolates, contributing to their genetic and phenotypic diversity (10, 11). For these reasons, *H. werneckii* is an emerging model for understanding eukaryotic adaptation to hypersaline environments and how hybridization events contribute to fungal adaptation to extreme environments (9).

Most of our understanding of *H. werneckii* biology and adaptation to hypersaline conditions have been derived from genomics, metabolic, and physiological analyses(1, 9–22). However, the lack of reliable genetic tools has limited the ability to test mechanistic hypotheses related to *H. werneckii* physiology and cell biology. Different methods exist to genetically transform fungi, including protoplast-mediated, *Agrobacterium*-mediate, electroporation, and biolistic(23, 24). These strategies require selectable markers, usually a drug-resistance gene or nutritional gene, to isolate cells that contain the desired construct. Recently, *in vitro* assembled CRISPR/Cas9 transformation systems have been utilized for highly efficient genetic manipulation of fungi (25–30). Due to its high efficiency, the CRISPR/Cas9 system has been adapted as a “marker-free” system to perform genome editing without needing a selectable marker (27).

Here, we adapted a protoplast-mediated transformation protocol to integrate a plasmid into the genome of *H. werneckii* ectopically. This ectopic integration strategy can be used to express the green fluorescent protein (GFP) in the cytoplasm of *H. werneckii*. We also develop a marker-free CRISPR/Cas9 protocol for knocking out genes in *H. werneckii*. These new strategies will further our understanding of the biology of this remarkable yeast yeast.

## Materials and Methods

### Strains, media, culture conditions

*H. werneckii* EXF-2000 was use for these studies. Yeast cells were grown in Glucose Minimal Media (GMM [Dextrose 10g/L, Trace Elements, Salt Solution]). Cells were grown at 30°C for 5 days unless otherwise specified.

### Protoplast generation

*Hortaea werneckii* cells were inoculated in GMM agar plates and incubated at 37°C for 5-7 days. Cells were harvested and washed once with osmotic media (1.2 M magnesium sulfate, 10 mM sodium phosphate buffer, pH 5.8). Digest the *H. werneckii* cell wall using osmotic media with 200 mg of Vinotaste for 4-8 hours at 30°C and shaking at 75 rpm. After digestion, two tubes were combined and overlayed with trapping buffer (0.6M sorbitol, 0.1 M Tris-HCl pH 7). Tubes then were centrifuged for 15 minutes at 2,500 g at 4°C. The cloudy layer (protoplast) at the interface of the osmotic media and trapping buffer was carefully moved into a new 15 mL conical and washed once with ice-cold STC (1.2M sorbitol, 10 mM CaCl_2_, 10 mM Tris-HCl pH 7.5). Protoplasts were resuspended with 1 mL of STC and counted using a hemocytometer.

### PEG-CaCl_2_-mediated transformation of pUCGH plasmid

200 µL of protoplast was transferred into a 1.5 mL and incubated with 1-5 µg of the pUCGH plasmid for 50 minutes to 1 hour. 1.25 mL of PEG-CaCl_2_ was added to the protoplast and incubated for 20 minutes. STC was added to protoplast up to 3 mL, and 300 µL were plated to SMM agar plates. Protoplasts were allowed to recover for 24 hours at room temperature. After 24 hours, the protoplast plates were overlayed with SMM top agar with 450 µg/ml of hygromycin B. Colonies appeared between 7-15 days of incubation. Individual colonies were transferred into fresh GMM agar with hygromycin B (150 µg/mL) and incubated at 30°C.

### Imaging EXF-2000

A wet mount of each strain was made by smearing *H. werneckii* cells in 20 µl of distilled water. Cells were then imaged using a widefield microscope (Leica DMi8) using a 100 X oil apochromat objective, and images were captured using a Leica K5 microscope camera.

### Assembly of CRISPR/Cas9 RNPs

5 µL of one crRNA (Table 1), tracrRNA, and nuclease-free duplex buffer (Integrated DNA Technology, IDT) were mixed and heated at 95°C for 5 minutes. Then, it was cool-down at room temperature to assemble the gRNA. 6 µL of each 33 µM gRNA, 6 µL of Cas9 nuclease (1µg/µL) (Integrated DNA Technology, IDT), and 8.5 µl of Cas9 working buffer were mixed and incubated for 5 minutes at room temperature. CRISPR/Cas9 complex was osmotically stabilized by adding an equal volume of 2X STC.

**Table 1:**
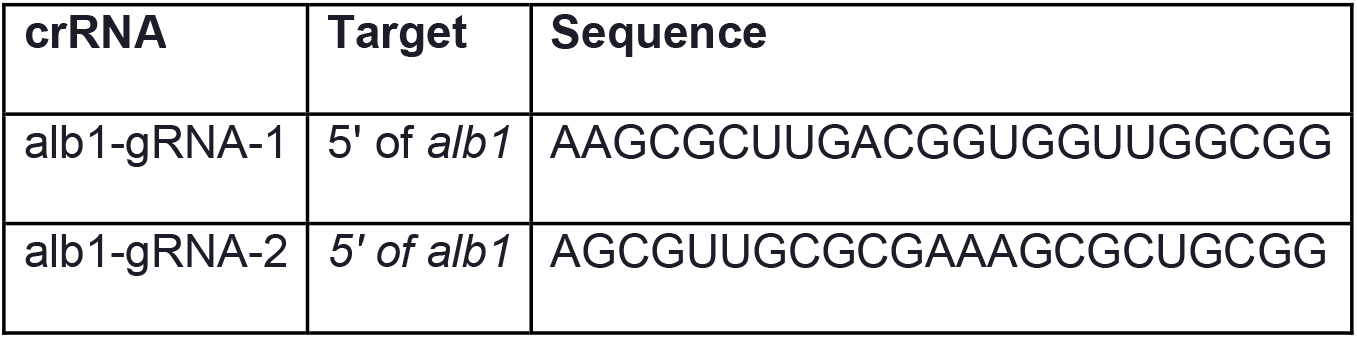
crRNAs used in this study.

### Marker-free CRISPR/Cas9-mediated transformation

Osmotically stabilized CRISPR/Cas9 complex, 70µL of protoplast, and 200 µl of PEG-CaCl_2_ were gently mixed in a 50 ml conical tube. STC replacing the osmotically stabilized CRISPR/Cas9 complex was used as a negative control. The protoplast mixture was incubated on ice for at least 30 minutes. 1 ml of PEG-CaCl_2_ was added and incubated for 15 minutes at room temperature. 325 µl of the protoplast were inoculated into 3 large SMM agar plates (150mm x 15mm) and incubated overnight at room temperature. Then, plates were incubated at 30°C until colonies started appearing on the plates.

### *PCR and sequencing of* alb1a *and* alb1b *locus*

200 mg of yeast cells were harvested from GMM plates and beadbeated for three 60s beadbeating and 30s resting cycles. Beadbeated cells were resuspended in 500µl CTAB, 20µl RNAase A, and 40 µl of Proteinase K. Resuspended cells were in incubated for 1 hour at 65°C. After incubation, cells were spun down for 10 minutes at 16,000g. DNA was purified using Promega MAXWELL RSC PureFood GMO and Authentication Kit following manufacture protocol. ProNex Size Selective Purification was used to remove the occasional melanin carryover. Paralog-specific primers were designed to amplify and sequence the *alb1a* or *alb1b* genes (Table 2). Amplicons were run in 0.8% agarose gels for 45 min at 150 V. Bands were cut and purified. Purified amplicons were sent to the Virginia Tech Genomics Sequencing Center for Sanger sequencing. Sequences were aligned using local MAFFTv7(31).

## Results

### Polyethylene glycol (PEG) calcium chloride protoplast-mediated transformation allows for ectopic integration of the pUCGH plasmid

*H. werneckii* susceptibility to drugs is determined by the environment (3, 4). We noticed that *H. werneckii* is susceptible to hygromycin B when grown on glucose minimum media, the same media commonly used to grow *Aspergillus fumigatus* (28). Based on this, we used hygromycin B as our selectable marker and used a sorbitol-stabilized glucose minimum media (SMM) to perform our transformations. We consistently obtained protoplast after 4-8 hours of digesting the cell wall using vinotaste (Novozymes). We then transformed the protoplast using a pUCGH vector, which contains the hygromycin B resistant gene (*hph*) under the control of *Aspergillus nidulans gpdA* promoter and eGFP under the control of the *Aspergillus oryzae* TEF1 promoter (32). We used a high quantity of plasmid to induce ectopic integration (5-15µg) of the plasmid. Protoplasts were allowed to recover for 24 hours before overlaying them with 10 ml of SMM top agar containing hygromycin B (450µg/ml). Hygromycin B-resistant colonies emerged after 7 days of incubation at 30°C (Fig 1A). Colonies were picked and streaked into a small petri dish containing GMM supplemented with hygromycin B (150µg/ml). Transformants expressed eGFP even after 5 passages in GMM+hygromycin B (Fig 1B). Thus, this approach can be utilized for ectopic integration of markers for cell biology studies.

**Figure 1.**
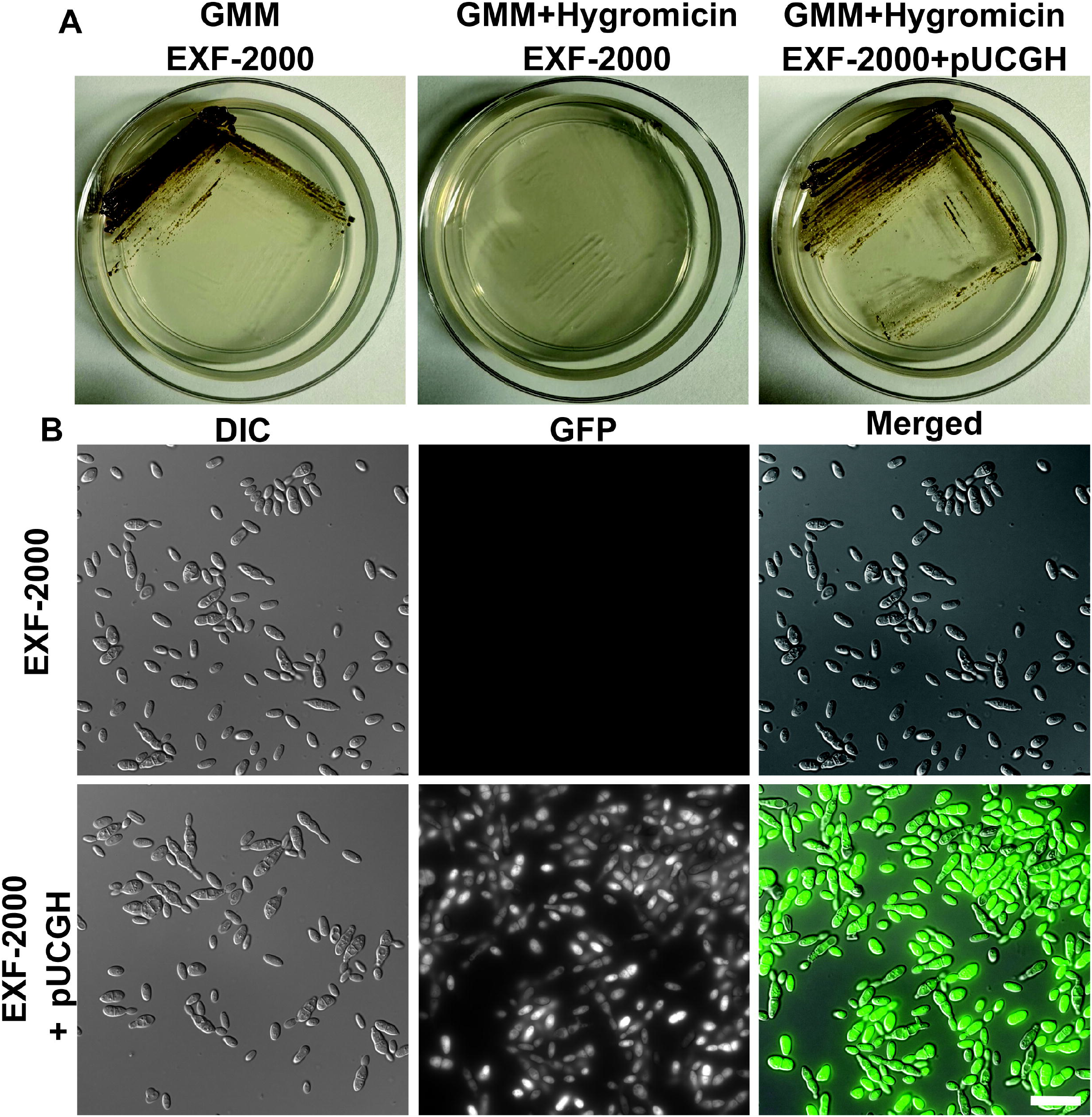
Integration of the pUCGH plasmid in *H. werneckii*. **(A)** *H. werneckii* EXF-200 is susceptible to hygromycin in GMM media, and hygromycin allows for the selection of ectopic integration of the pUCGH plasmid. Plates were streaked into the respective media, incubated at 37°C, and photographed after 5 days. **(B)** *H. werneckii* with integrated pUCGH expresses eGFP under the *A. oryzae* tef1 promoter. Micrographs were obtained using a 100X objective. Scale bar, 20 µm.

### CRISPR/Cas9 marker-free transformation

More targeted genetic approaches are needed to further our understanding of *H. werneckii* biology. Similar to other ascomycetes, adapting the PEG-CaCl2 mediated transformation protocol has proven challenging for targeted gene manipulation due to the low homologous recombination rate (33). This challenge is further exacerbated due to the EXF-2000 strain having a significant portion of its genome duplicated due to an intraspecific hybridization event (10). Due to this challenge, we decided to adopt a CRISPR/Cas9 marker-free approach and target the *H. werneckii* melanin synthesis pathway to have a visual phenotype we could screen(27). We identified 2 copies of the alb1 gene–*alb1a* (BTJ68_00107) and *alb1b* (BTJ68_01291)–using FungiDB(34). We targeted a conserved region between *H. werneckii’s* two *alb1* paralogs: using 2 gRNAs (Table 1). We generated protoplast using the same approach for the ectopic integration, and without standardizing the number of protoplasts, we obtained approximately 3-5 colonies that exhibited an albino phenotype (Fig. 2A) per transformation. We decided to determine if the concentration of protoplasts might impact the efficiency of our CRISPR/Cas9 approach (Fig. 2B). We observed that transforming a mix containing 10^4^ protoplasts led to approximately 6.5% success rate, compared to the ∼1.9% success rate when transforming 10^5^ protoplasts. We did not observe any spontaneous albino mutant arising from the protoplast generation or the exposure of the protoplast to STC. The albino mutants could still not produce melanin after 5 passages, indicating that the deletion of the *alb1* genes was stable (Fig. 2C). These albino mutants retain the same cell morphology as the wild-type when grown in GMM (Fig 2D), indicating that melanin does not impact cell morphology in these conditions. A 1.5 kb genomic region that covers the site targeted by CRISPR/Cas9 was amplified using primers specific to each *alb1* paralog (Table S1). Gel electrophoresis for the *alb1a* PCR reaction showed bands of similar size between the three independent mutants and the wild-type EXF-2000 strain, ∼1.5 kb (Fig 3A). Similarly, the Δ*alb1*.*1* and Δ*alb*.*3* had *alb1b* amplicons of comparable size to the EXF-2000 band (Fig. 3B). In contrast, the Δ*alb1*.*2* had a smaller amplicon of ∼1kb, and this mutant had a deletion in their *alb1b* gene of ∼500bp. In the *alb1a* locus, we found that all the Δ*alb1*strains had similar mutations constrained between the Cas9 cut sites (Fig 3C). The Δ*alb1*.*1* and Δ*alb1*.*3* followed a similar pattern for the *alb1b* locus (Fig.3D). In concordance with the amplicon size, the Δ*alb1*.*2 alb1b* locus had a 528 bp deletion. The EXF-2000 *alb1a* and *alb1b* loci sequences are in the reference genome.

**Figure 2.**
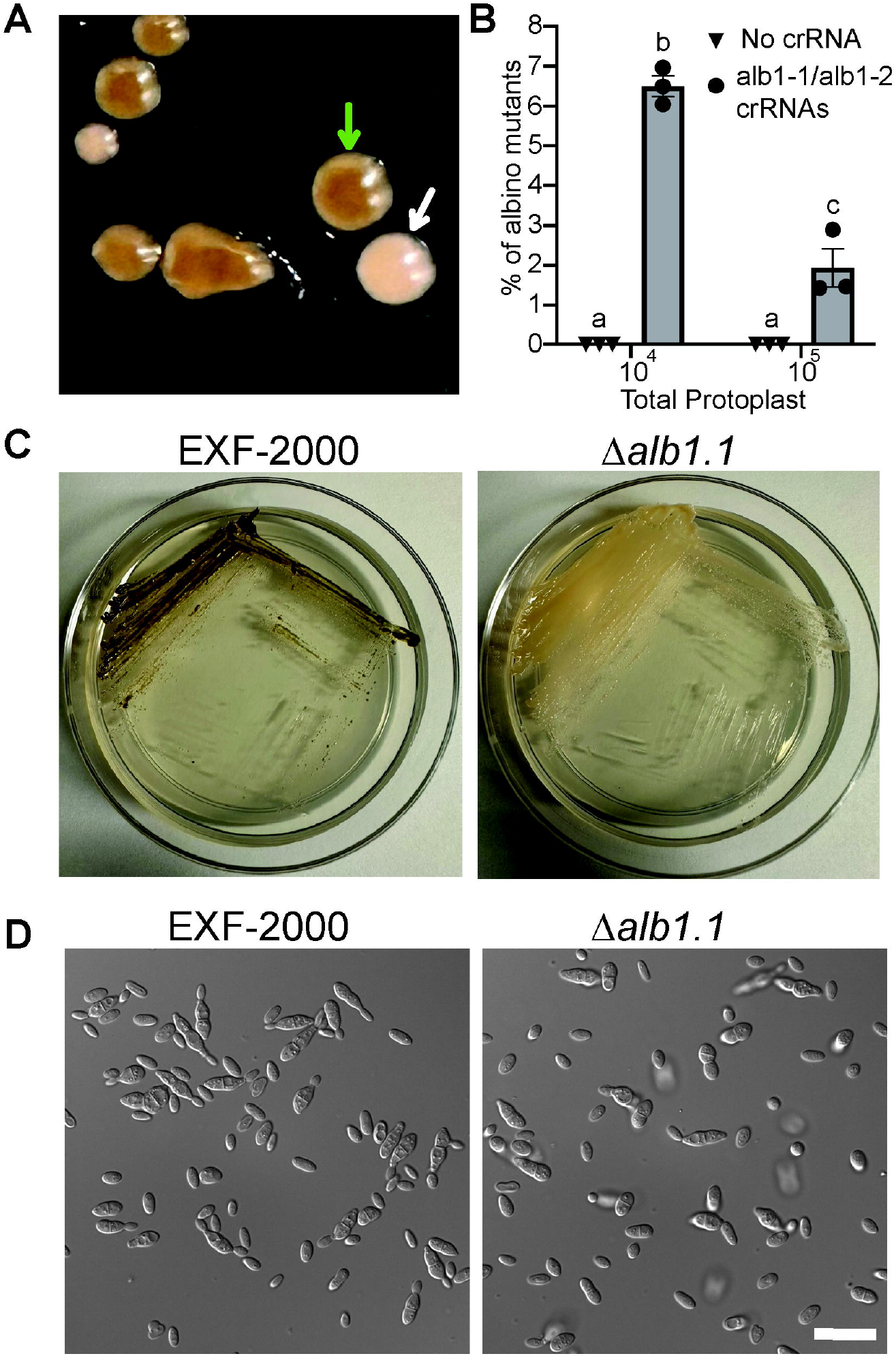
Generation of albino mutants using CRISPR/Cas9. **(A)** Albino mutant in transformation plate (white arrow) next to melanized EXF-2000 (green arrow) **(B)** A lower amount of protoplast increases the efficiency of CRISPR/Cas9 transformation. 10^4^ or 10^5^ protoplasts were transformed with the CRISPR/Cas9 complex with each crRNA (circles) or STC (inverted triangle). STC-only protoplasts did not result in spontaneous albino mutants. Using 10^4^ protoplasts led to an average of 6.5% albino mutants, in contrast to the 1.9% when using 10^5^ protoplasts. 2-way ANOVA followed by Tukey’s post hoc test to compare the means of each experimental condition. Statistical significance was determined by p≤0.05 **(C)**. The Δ*alb1*.*1* fails to melanize after prolonged incubation in GMM. Plates were streaked and incubated at 30°C for 5 days before being photographed. **(D)** Δ*alb1*.*1* has a similar morphology as the parent EXF-2000 strain when grown in GMM agar. Micrographs were obtained using a 100X objective. Scale bar, 20 µm.

**Figure 3.**
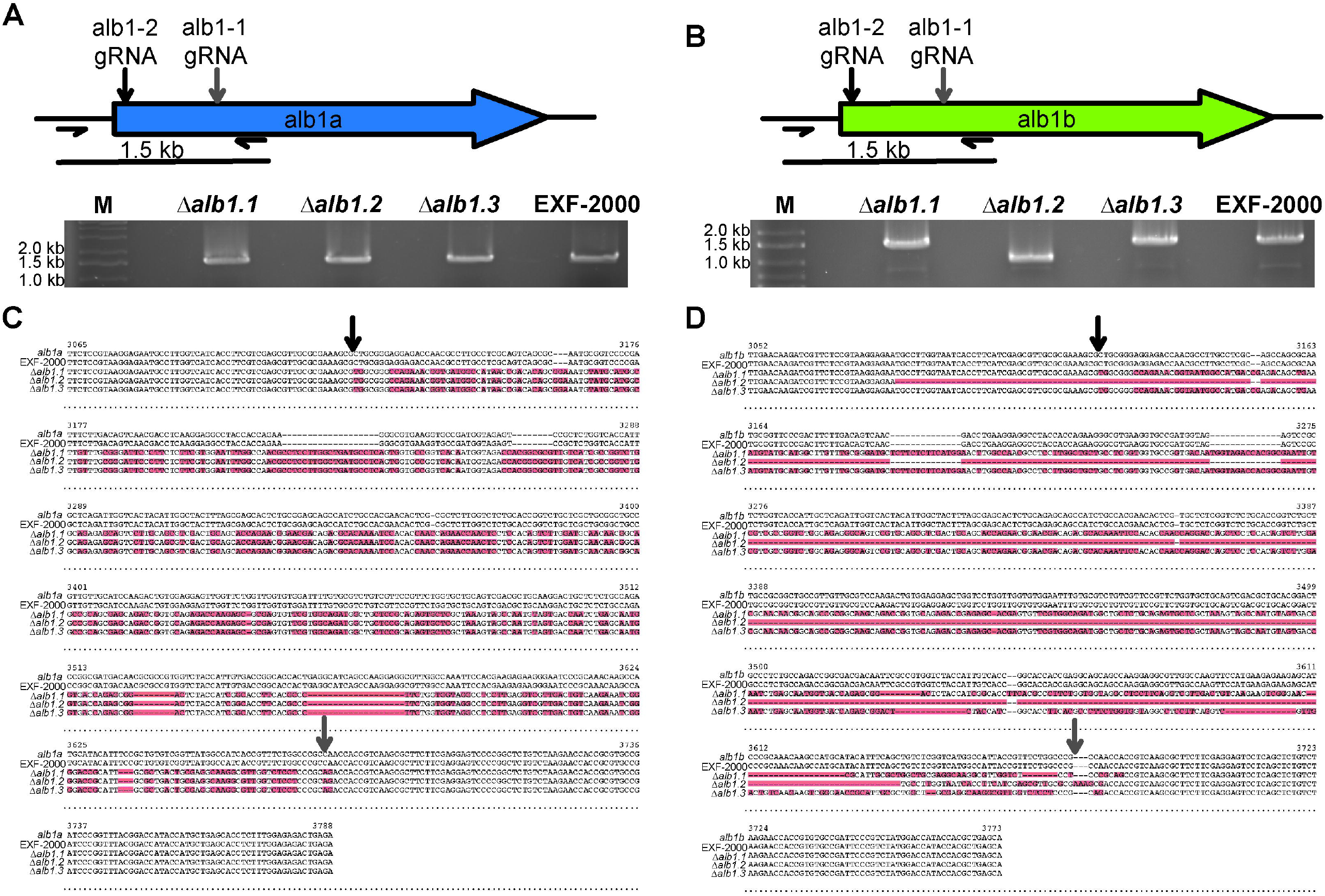
*alb1a* and *alb1b* loci are mutated in the Δ*alb1* strains. **(A-B)** Gel electrophoresis of the PCR amplicons of the *alb1a* **(A)** and *alb1b* **(B)** 5’ gene region. M, marker (GeneRuler 1KB plus). **(C-D)** Alignment of the *alb1a* **(C)** and *alb1b* **(D)** loci. The gray arrow points at the expected cut site for Cas9 loaded with alb1-1 gRNA, and the black arrow points at the predicted cut site for Cas9 loaded with alb1-2 gRNA. Red highlights point to areas of divergence between the sequence and the reference genome (*alb1a* or *alb1b*).

## Discussion

*Hortaea werneckii* has a unique cell cycle division and can grow at high salt concentrations (3, 4, 35). Due to most isolates being intraspecific hybrids with significant phenotypic variation, *H. werneckii* can become a model for understanding how hybridization leads to novel phenotypes (3, 9). However, the lack of genetic tools has limited the ability to study this organism. Here, we develop two different genetic transformation approaches to further our studies of this unconventional yeast. First, we have developed a plasmid-based ectopic integration protocol. This system can be used to express cellular markers for live cell imaging. For example, we could use it to insert a histone-GFP plasmid and monitor nuclear dynamics. Second, we develop a marker-free CRISPR/Cas9 protocol to target specific genes.

One caveat is that our current CRISPR/Cas9 protocol still needs to be more effective and would not be practical for genes that would not have a visual readout. On average, we had a 6-7% success rate on targeting both gene paralogs, while other organisms had a 50% to 90% success rate (27–29). However, these organisms were haploid, and only one copy of the gene needed to be mutated, not the 2 copies of most genes in *H. werneckii*. Lastly, as there is a marker system and CRISPR/Ca9 can overcome the homologous recombination rate, it is possible to combine both methods to develop a marker-based CRISPR/Cas9 approach and select those mutants with successful integration of our cassette of interest (28). At the moment of this publication, only one marker has been developed (hygromycin) for *H. werneckii*. Thus, this approach might only be limited to protein tagging or deletion of genes with only 1 copy. Nonetheless, other drugs can be explored to expand the available range of selectable markers.

## Acknowledgments

The authors thank Dr. William Steinbach and Dr. Praveen Juvaadi for sharing the pUCGH plasmid. We also want to thank Dr. Rachel Whitaker, Dr. Scott Dawson, and the Marine Biological Laboratory’s (MBL) Microbial Diversity course for allowing us to use their microscope and resources. The author wants to thank Norman van Rhijn (Manchester Fungal Infection Group) and Robb Cramer’s Lab (Dartmouth) for sharing their CRISPR/Cas9 protocols that were the basis for developing our protocol. This project was supported through the L. & A. Colwin Summer Research Fellowship awarded to JVM. JVM wants to thank William Wolf (Virginia Tech), and the Vargas-Muñiz Lab for critically reading this manuscript. Lastly, we want to recognize FungiDB and VEuPathDB, which are invaluable resources for the community, and this project would not have been possible without their support and tools.

## Figure Legends

**Table S1:**
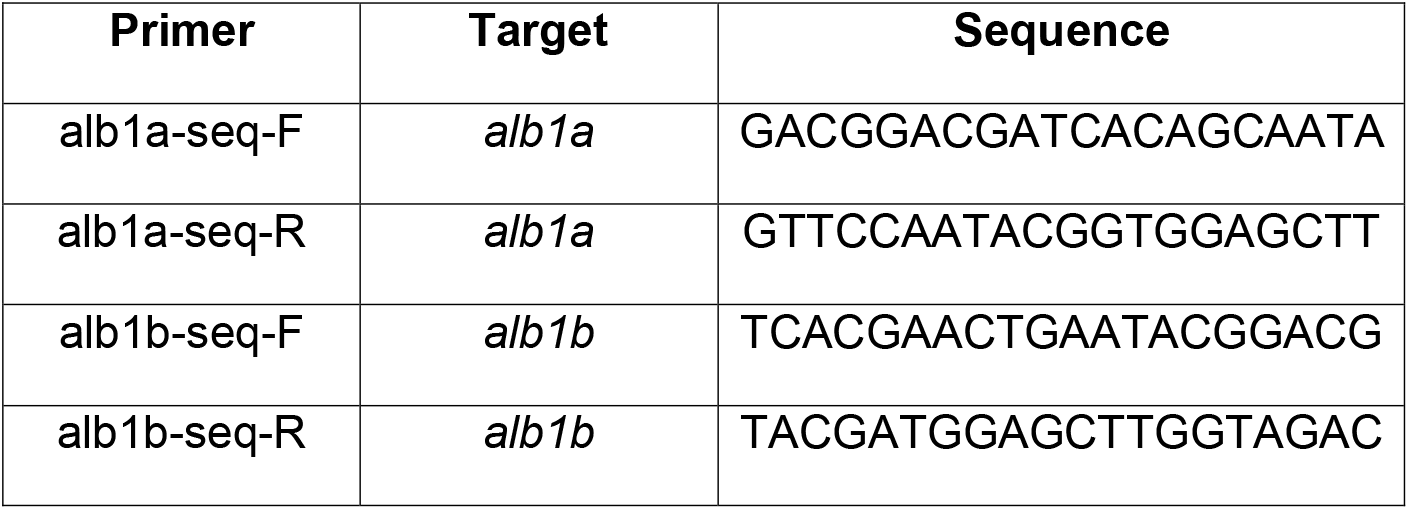
Primers used in this study.

## Notes

### Competing Interest Statement

The authors have declared no competing interest.

### Summary of Updates

We sequenced our mutants and also determined if the number of protoplast impacts the efficacy of transformation. Fig 2 was revised to include some of our findings, and we added a new Figure 3 to include the sequencing of the albino mutants.

